# Charting the native architecture of thylakoid membranes with single-molecule precision

**DOI:** 10.1101/759001

**Authors:** Wojciech Wietrzynski, Miroslava Schaffer, Dimitry Tegunov, Sahradha Albert, Atsuko Kanazawa, Jürgen M. Plitzko, Wolfgang Baumeister, Benjamin D. Engel

## Abstract

Thylakoid membranes scaffold an assortment of large protein complexes that work together to harness the energy of light to produce oxygen, NADPH, and ATP. It has been a longstanding challenge to visualize how the intricate thylakoid network organizes these protein complexes to finely tune the photosynthetic reactions. Using cryo-electron tomography to analyze membrane surface topology, we have mapped the native molecular landscape of thylakoid membranes within green algae cells. Our tomograms provide insights into the molecular forces that drive thylakoid stacking and reveal that photosystems I and II are strictly segregated at the borders between appressed and non-appressed membrane domains. This new approach to charting thylakoid topology lays the foundation for dissecting photosynthetic regulation at the level of single protein complexes within the cell.

---

Membranes orchestrate cellular life. In addition to compartmentalizing the cell into organelles, membranes can organize their embedded proteins into specialized domains, concentrating molecular partners together to drive biological processes *(1, 2)*. Due to the labile and often transient nature of these membrane domains, it remains a challenge to study how individual protein complexes are organized within them.

The question of membrane domain organization is especially pertinent to thylakoids, sheet-like membrane-bound compartments that produce oxygen while converting light energy into biochemical energy, thereby sustaining most of the life on Earth. These light-dependent photosynthetic reactions are driven by the concerted actions of four large protein complexes within the thylakoid membrane. Photosystem II (PSII), cytochrome *b*_*6*_*f* (cyt*b*_*6*_*f*), and photosystem I (PSI) form an electron transport chain that produces NADPH and pumps protons into the thylakoid lumen. ATP synthase then uses this electrochemical gradient across the membrane to generate ATP. In most photosynthetic eukaryotes, the thylakoid membranes are subdivided into appressed regions that face adjacent membranes within thylakoid stacks (called grana in higher plants) and non-appressed regions that freely face the stroma (*3–6*). In both plants and algae, PSII and PSI appear to be segregated to the appressed and non-appressed membranes, respectively (*6–9*). This lateral heterogeneity is believed to coordinate photosynthesis by concentrating different reactions within specialized membrane domains, while the redistribution of light-harvesting complex II (LHCII) antennas between these domains may enable adaptation to changing environmental conditions *(10, 11)*.

Much of what is known about the molecular organization of thylakoids comes from freeze-fracture electron microscopy (and the related deep etch technique), which provides views of membrane-embedded protein complexes within the cell *(12)*. Combined with biochemical fractionation *(13, 14)*, these membrane panoramas helped describe the lateral heterogeneity of thylakoids *(7, 8, 15)*. However, the platinum replicas produced by this technique have limited resolution and only provide access to random fracture planes through the membranes. More recently, atomic force microscopy (AFM) has been used to map the macromolecular organization of thylakoids *(16, 17)*. AFM can very precisely measure topology and thus, can distinguish between each of the major photosynthetic complexes within a hydrated membrane. However, only one membrane surface can be visualized at a time, and the membranes must first be removed from the cell using detergents. As an alternative, cryo-electron tomography (cryo-ET) has been shown to resolve PSII complexes within hydrated thylakoids *(18–20)*. Although multiple overlapping membranes can be imaged with this technique, the thylakoids in these studies were isolated from the chloroplast in order to produce sufficiently thin samples.

Dissecting the interrelationship between membrane domains and thylakoid architecture requires a molecular view of intact thylakoid networks within native cells. In pursuit of this goal, we combined cryo-focused ion beam milling *(21, 22)* with cryo-ET *(23)* to image thylakoid membranes within vitreously-frozen *Chlamydomonas reinhardtii* cells. Several years ago, we demonstrated how this approach can capture the undisturbed membrane architecture of this green alga, revealing an elaborate system of stacked thylakoids *(4)*. However, due to the limited resolution of the CCD cameras used at that time, this investigation was limited to a description of membrane architecture, without visualizing the protein complexes embedded within these membranes. Here, we leverage advances in direct detector cameras and the contrast-enhancing Volta phase plate *(24)* to resolve the thylakoid-embedded complexes, enabling us to describe the molecular organization of thylakoids *in situ*, within their native cellular context.

Our tomograms show numerous densities corresponding to ribosomes and photosynthetic complexes decorating the appressed and non-appressed surfaces of the thylakoid network (Figs. 1A-B, S1, Movie S1). Close visual inspection revealed the unmistakable shapes of ATP synthase and PSII protruding into the stroma and lumen, respectively (Fig. 1C-D). In order to map these decorating densities onto the native thylakoid architecture, we developed a visualization approach called a “membranogram”, where densities from the tomogram are projected onto the surface of a segmented membrane (Fig. 1E). The result is a topological view of the membrane surface that resembles AFM data. However, unlike AFM, membranograms can display the topology of both sides of each membrane within the cellular volume. By dynamically growing and shrinking the segmentations, the membranograms allow us to track how densities change as they extend from the membrane surface and compare these densities to mapped in molecular models of different complexes (Movie S2). This enabled the manual assignment of thylakoid-associated complexes by considering whether the densities projected into the stroma or thylakoid lumen (Fig. 2A) and then comparing the densities to known structures of each complex (Fig. S2). We used manually-picked positions to generate subtomogram averages of PSII and ATP synthase, structurally confirming that membranograms can correctly identify these complexes (Fig. S3).

**Figure 1.**
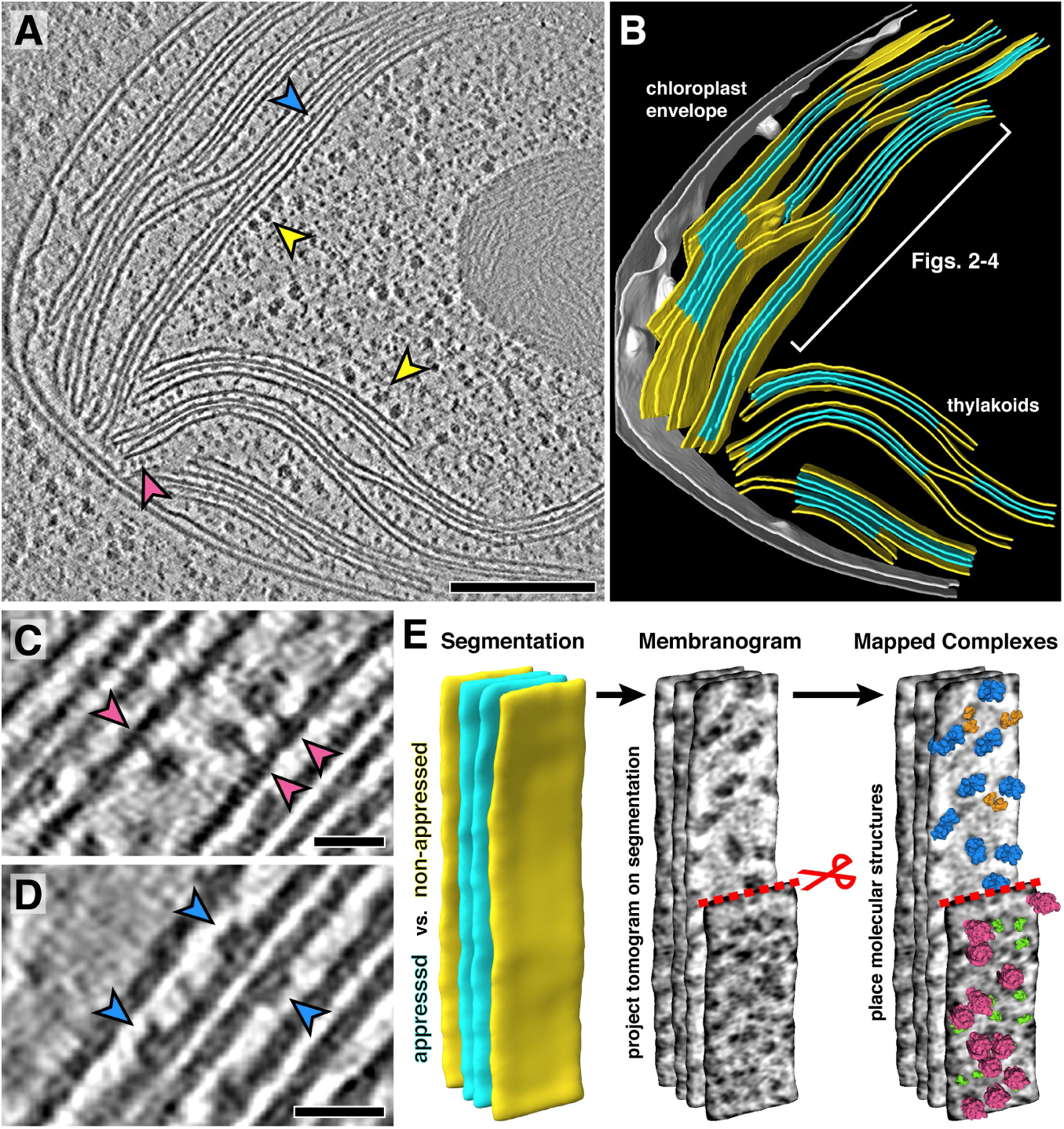
In situ cryo-electron tomography reveals the native molecular architecture of thylakoid membranes. **A)** Slice through a tomogram of the chloroplast within an intact Chlamydomonas cell. Arrowheads point to membrane-bound ribosomes (yellow), ATP synthase (magenta), and PSII (blue). **B)** Corresponding 3D segmentation of the chloroplast volume, with non-appressed stroma-facing membranes in yellow, appressed stacked membranes in blue, and the chloroplast envelope in grey. The thylakoid region used in Figs. 2-4 is indicated. **C-D)** Close-up views, showing individual ATP synthase (C) and PSII (D) complexes. **E)** Mapping photosynthetic complexes onto thylakoids using membranograms. Segmented membranes are extracted from the tomogram (left). Tomogram densities are projected onto the segmented surfaces, showing densities that protrude from the membranes (middle, red scissors: part of the non-appressed membrane has been removed to reveal luminal densities on the appressed membrane). Protein complexes are mapped onto membranograms based on the shapes of the densities and whether they protrude into the stromal or luminal space (blue: PSII, orange: cyt*b*_*6*_*f*, green: PSI, magenta: ATP synthase). Scale bars: 200 nm in A, 20 nm in C and D. See Movies S1 and S2.

**Figure 2.**
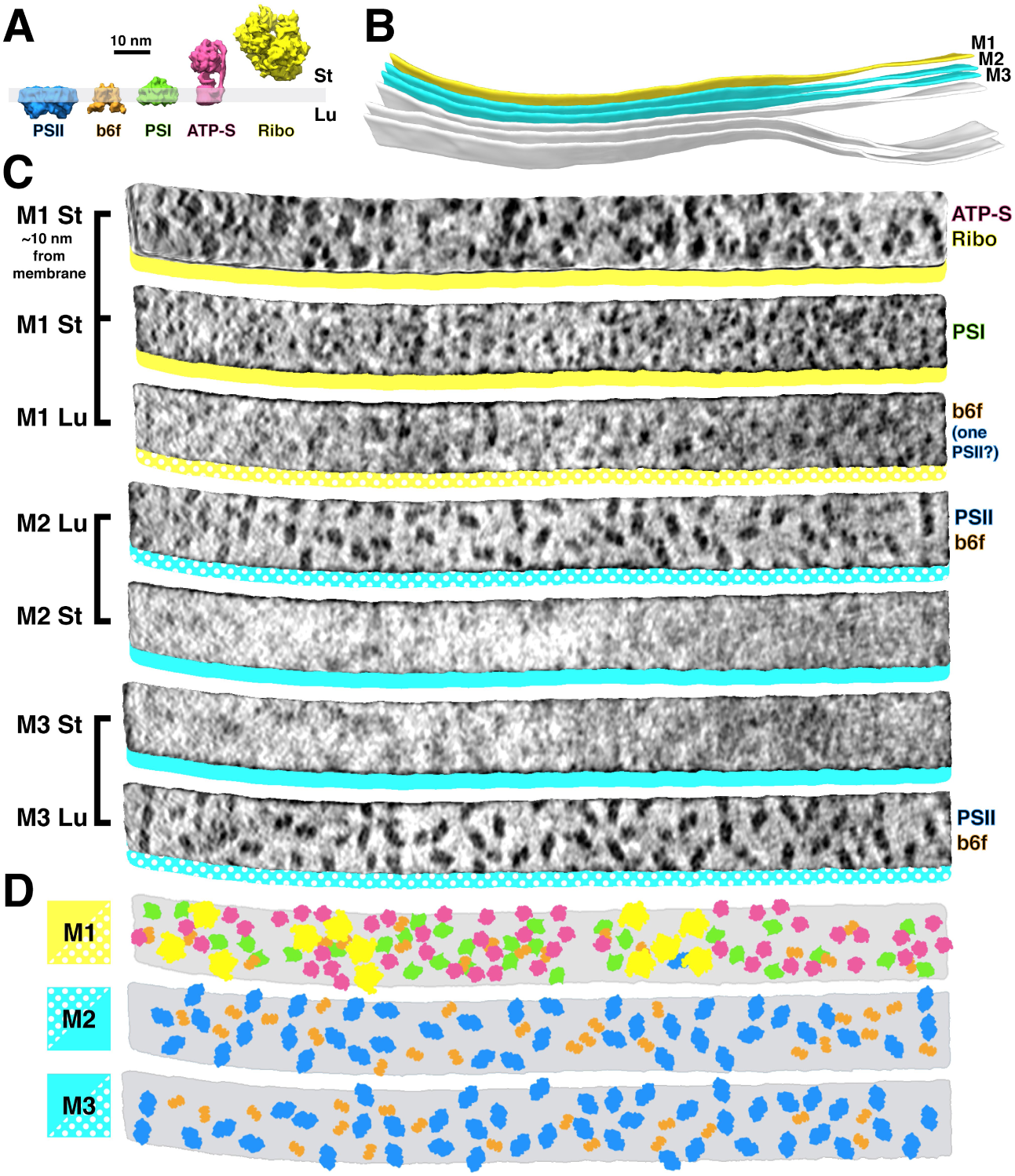
Mapping photosynthetic complexes within appressed and non-appressed thylakoid membranes. **A)** Schematic of how each molecular complex extends from the membrane into the stroma (St) and thylakoid lumen (Lu). Into the lumen, large dimeric densities extend ∼ 4 nm from PSII and small dimeric densities extend ∼ 3 nm from cyt*b*_*6*_*f* (b6f). Into the stroma, a small monomeric density extends ∼ 3 nm from PSI, the F_1_ region of ATP synthase (ATP-S) extends ∼ 15 nm, and thylakoid-bound ribosomes (Ribo) extend ∼ 25 nm. **B)** Three segmented thylakoids from the region indicated in Fig. 1B. Membranes 1-3 (M1-M3, yellow: non-appressed, blue: appressed) are examined by membranograms. **C)** Membranogram renderings of M1-M3. All membranograms show the densities ∼ 2 nm above the membrane surface, except the top membranogram, which was grown to display densities ∼ 10 nm into the stroma. Stromal surfaces are underlined with solid colors, whereas luminal surfaces are underlined with a dotted color pattern. The complexes identified in each surface are indicated on the right. **D)** Model representation of M1-M3, showing the organization of all of the thylakoid complexes (colors correspond to the schematic in A). See Fig. S2.

We used membranograms to examine the molecular organization within appressed and non-appressed domains of the thylakoid network (Fig. 2). The complexes in appressed membranes (M2 and M3 in Fig. 2B-D) could be reliably assigned. The stromal surfaces of these membranes showed no large densities, whereas two clearly distinguished classes of densities were seen on luminal surfaces: large PSII dimers and smaller cyt*b*_*6*_*f* dimers that extend ∼ 4 nm and ∼ 3 nm into the lumen, respectively. Non-appressed membranes (M1 in Fig. 2B-D) were more complicated to analyze. We first assigned thylakoid-associated ribosomes and ATP synthases with high certainty by growing the segmentation ∼ 10 nm away from the membrane. Assignment of PSI was more difficult: we marked single densities that protruded ∼ 3 nm into the stroma and were not directly under a ribosome or ATP synthase. On the luminal side, almost no large densities were observed that could correspond to PSII. Although we did see frequent small dimers that resembled cyt*b*_*6*_*f*, they were more difficult to identify due to increased background signal on the luminal surfaces of non-appressed membranes. We generated membrane models from the assigned particles (Fig. 2D) to analyze how the complexes are arranged within the plane of the membrane. In total, we quantified 84 membrane regions from four tomograms (Table 1) and found clear evidence of lateral heterogeneity: PSII was almost exclusively found in the appressed regions, whereas PSI, ATP synthase, and ribosomes were restricted to the non-appressed regions. Cyt*b*_*6*_*f* was observed with almost equal abundance in both regions. Clustered poly-ribosome chains were clearly resolved decorating the non-appressed thylakoids (Figs. 1A, 2D-C). In contrast, PSI, PSII, cyt*b*_*6*_*f*, and ATP synthase were distributed relatively evenly along their respective membrane regions (nearest-neighbor distances plotted in Fig. S4). The concentrations we measured for different photosynthetic complexes were similar to previous biochemical estimates *(14, 25, 26)*, as well as AFM measurements *(17)* and counts of “membrane-embedded particles” from freeze-fracture *(8)*, indicating that direct comparisons can be made between cryo-ET data and these earlier studies.

**Table 1.** Average concentrations of macromolecular complexes per square micrometer of native thylakoid membranes. N= 84 membrane regions (51 non-appressed and 33 appressed) from four tomograms. “Unknown” densities are particles that could not be assigned an identity, including particles found near the edges of the segmented membranes. Asterisks show classes of complexes that were identified with lower confidence.

|  | PSII | *Cytb<sub>6</sub>f* | PSI | ATP-S | Ribo | Unknown | Total |
| --- | --- | --- | --- | --- | --- | --- | --- |
| Non-appressed | 24 | 501* | 1049* | 1652 | 113 | 1568 | 4907 |
| Appressed | 1122 | 631 | 2 | 0 | 0 | 170 | 1925 |

At the boundary between appressed and non-appressed membranes, it is widely believed that there is a specialized domain called the “grana margin” where PSII and PSI intermix (14, 27–29). Using membranograms, we visualized the molecular organization of native thylakoids that transitioned between appressed and non-appressed regions (Figs. 3 and S5). Strikingly, we observed a sharp boundary between PSII and PSI that exactly matched the division in thylakoid architecture. The photosystems did not intermix, and we saw no clear difference in particle abundance near the boundary. We therefore find no evidence for grana margins in *Chlamydomonas* cells grown under these conditions (see methods).

**Figure 3.**
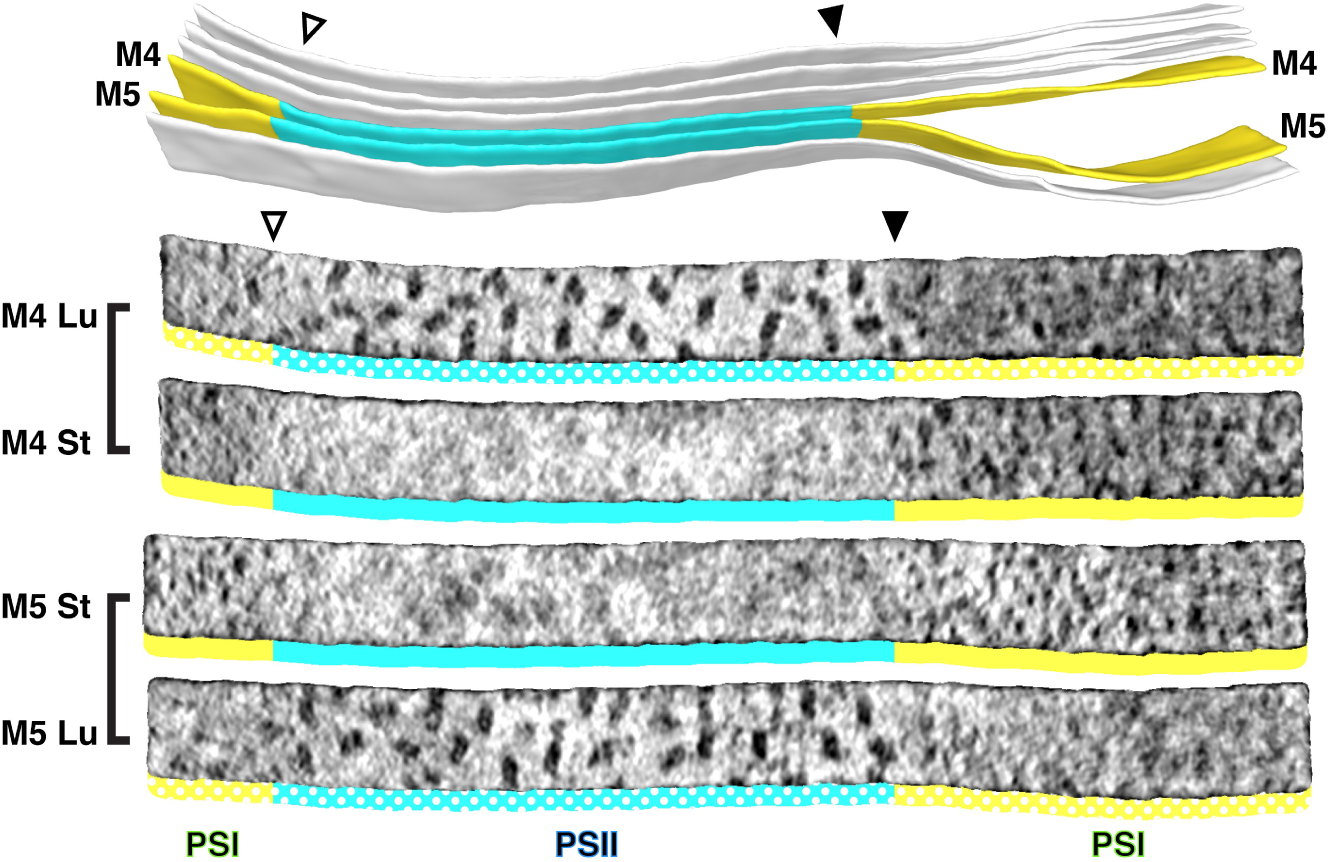
Strict segregation of PSII and PSI at transitions between appressed and non-appressed regions. **Above:** Three segmented thylakoids from the region indicated in Fig. 1B. Membranes 4-5 (M4 and M5, yellow: non-appressed, blue: appressed) are examined by membranograms. **Below:** Membranograms of M4 and M5. All membranograms show the densities ∼ 2 nm above the membrane surface. Stromal surfaces are underlined with solid colors, whereas luminal surfaces are underlined with a dotted color pattern. Transitions between appressed and non-appressed regions are marked with arrowheads. PSII is exclusively found in the appressed regions, whereas PSI is exclusively found in the non-appressed regions, with sharp partitioning at the transitions between regions. For an additional example of how lateral heterogeneity of PSII and PSI is coupled to membrane architecture, see Fig. S5.

What drives the strict lateral heterogeneity that we observe between appressed and non-appressed domains? PSI is presumably excluded from appressed membranes because its ∼ 3 nm stromal density is too bulky to fit into the ∼ 3 nm space between stacked thylakoids *(4, 18, 30)*. Conversely, PSII and its associated LHCII antennas may induce thylakoid stacking, a causal relationship that would precisely limit PSII to appressed membranes. Several studies have observed semi-crystalline arrays of C_2_S_2_-type *(18, 31)* or C_2_S_2_M_2_-type *(31)* PSII-LHCII super-complexes in isolated thylakoids, and it has been proposed that the overlap of LHCII or PSII between membranes mediates thylakoid stacking *(18, 31, 33, 34)*. Although we observed randomly oriented PSII complexes instead of ordered arrays, we nonetheless looked for evidence of supercomplex interactions across native thylakoid stacks (Fig. 4). We first created membranogram overlays of adjacent membranes spanning either the thylakoid lumen or stromal gap (Fig. 4B). Then we generated membrane models by using the positions and rotational orientations of PSII luminal densities seen in the membranograms to place structures of C_2_S_2_M_2_-type PSII-LHCII supercomplexes *(35)* (Fig. 4C-D). Interestingly, we observed almost no over-lap between the supercomplex models within the plane of the membrane (Fig. 4C), indicating that there is ample space with-in these appressed regions to accommodate large C_2_S_2_M_2_-type supercomplexes. The supercomplexes occupied ∼ 45% of the membrane surface area (Fig. 4E), with cyt*b*_*6*_*f* occupying an additional ∼ 5%. Thylakoids are ∼ 70% protein *(36)*, suggesting that other complexes such as extra LHCII antennas may occupy up to 20% of the surface area. This spacing should also allow room for rapid diffusion of plastoquinone between PSII and cyt*b*_*6*_*f* within appressed membranes.

**Figure 4.**
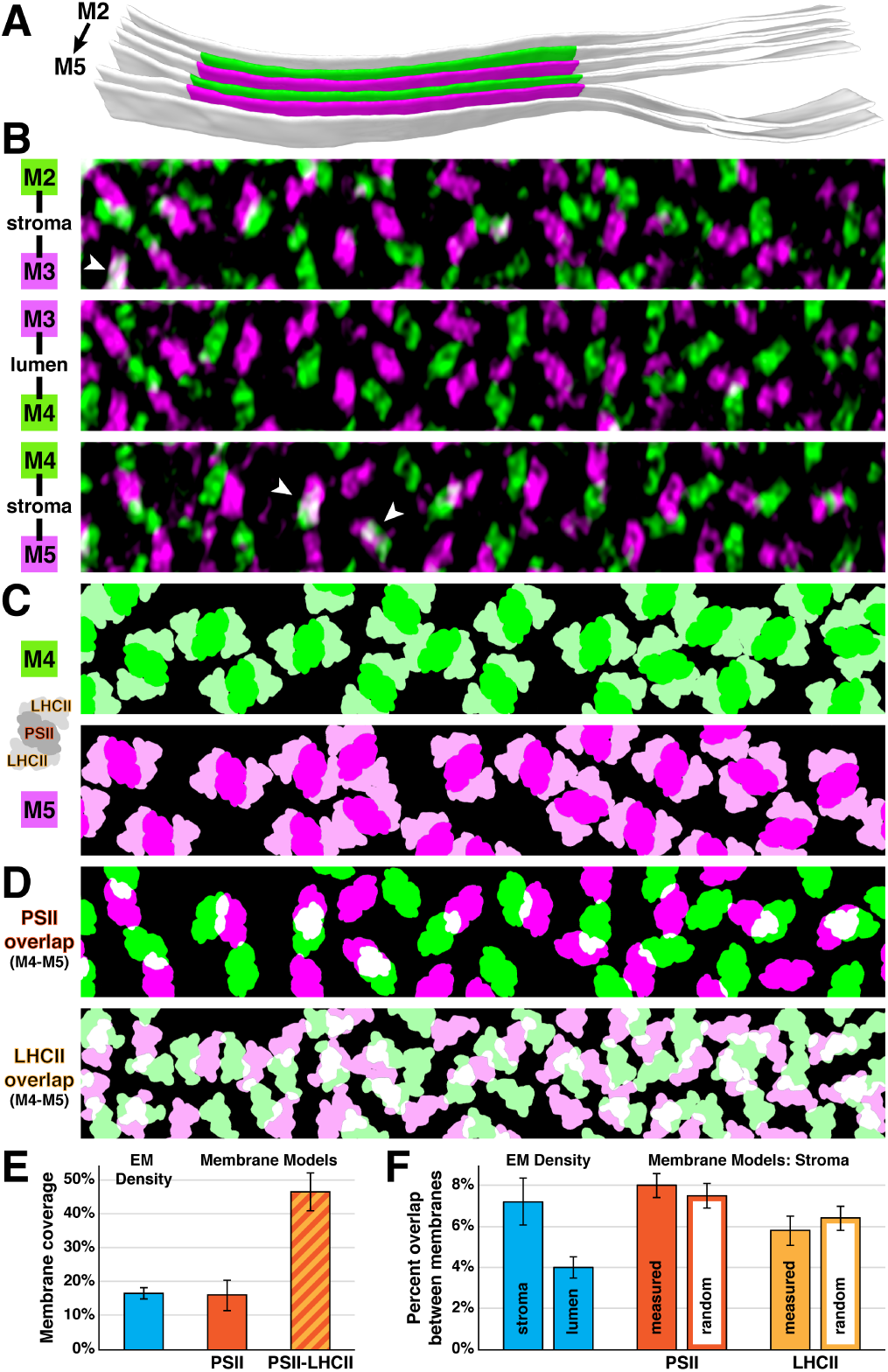
Native thylakoids can accommodate PSII-LHCII supercomplexes, which are randomly distributed across the stromal gap. **A)** Three segmented thylakoids from the region indicated in Fig. 1B. Membranes 2-5 (M2 to M5, alternating green and magenta) are examined by membranograms. **B)** Overlays of two luminal surface membranograms, superimposed across the stromal gap (M2-M3, M4-M5) and thylakoid lumen (M3-M4). Membranograms are pseudocolored corresponding to A. Overlapping PSII complexes are indicated with white arrowheads. **C)** Model representations of M4 and M5, with C_2_S_2_M_2_-type PSII-LHCII supercomplexes positioned according to the luminal densities observed in the membranograms. The spacing indicates that appressed thylakoids can accommodate these supercomplexes. **D)** M4-M5 overlays using the membrane models from C, separately showing the PSII core complexes (top) and LHCII antennas (bottom). Overlapping regions are in white. **E)** The percentage of membrane surface area occupied by luminal EM density (15.9% ± 2.0%, N= 19 membranes) in membranograms (e.g., panel B), and occupied by PSII core complexes (16.0% ± 4.5%, N= 9 membranes) or C_2_S_2_M_2_-type PSII-LHCII supercomplexes (46.4% ± 5.5%, N= 9 membranes) within membrane models (e.g., panel C). **F)** Blue bars: The percentage of luminal EM density in membranograms that overlaps between adjacent appressed membranes spanning the stromal gap (7.2% ± 2.3%, N= 11 overlays) and thylakoid lumen (4.0% ± 1.0%, N= 6 overlays). See Fig. S6 for how the EM density was thresholded to calculate surface area and overlap. Red and yellow bars: Using membrane models, the percentage of PSII and LHCII surface area that overlaps between adjacent appressed membranes spanning the stromal gap (N= 4 overlays). The experimental measurements (PSII: 8.0% ± 0.6%, LHCII: 5.8% ± 0.7%) and simulations of complexes randomly positioned within the membrane (PSII: 7.5% ± 0.6%, LHCII: 6.4% ± 0.6%, N= 100 simulations per overlay) were not significantly different (P > 0.05 from t-test with Welch’s correction for unequal variance: P= 0.174 for PSII, P= 0.197 for LHCII). See Fig. S7 and methods for how the random simulations were performed. Error bars denote standard deviation.

The membranogram overlays showed very little overlap of PSII between adjacent membranes. In particular, PSII complexes across the thylakoid lumen appeared to completely interdigitate, with only ∼ 4% overlap (Fig. 4B and F). PSII interdigitation has previously been observed in isolated pea thylakoids with a contracted ∼ 4.5 nm lumen, which sterically could only permit interdigitation. However, the thylakoids in our light-adapted *Chlamydomonas* cells have a ∼ 9 nm lumen, which is enough space to permit face-to-face interactions of PSII across the lumen, yet the PSII complexes remain interdigitated. It has been suggested that interdigitation could facilitate diffusion of the soluble electron carrier plastocyanin through the lumen, while also enabling the lumen to contract and block diffusion in response to changing light conditions *(30)*. To investigate interactions across the stromal gap, we measured the intermembrane overlap from the membrane models of C_2_S_2_M_2_-type supercomplexes and compared it to simulated data where the same number of supercomplexes were randomly positioned (Fig. 4D and F). We found that the overlap of PSII-to-PSII and LHCII-to-LHCII were both statistically indistinguishable from random. It remains to be tested whether randomly distributed interactions covering only ∼ 8% of the PSII surface area and ∼ 6% of the supercomplex LHCII surface area can mediate the tight adhesion between stacked thylakoid membranes.

*In situ* cryo-ET has enabled us to visualize the native architecture of thylakoids with molecular resolution, revealing several striking findings. We observe sharp boundaries between PSII and PSI that are tightly coupled to membrane architecture, with no evidence of grana margins at transitions between appressed and non-appressed membrane domains. Contrary to previous reports of crystalline PSII arrays in isolated spinach, pea, and diatom thylakoids *(18, 20, 31, 32, 37)*, we found that PSII complexes are randomly oriented within native appressed Chlamydomonas thylakoids, with ample spacing to accommodate C_2_S_2_M_2_-type supercomplexes. This observation supports the physiological relevance of these larger supercomplexes, which have been primarily characterized *in vitro (35, 38)*. Our analysis also raises questions about the forces that shape thylakoid architecture, challenging the idea that thylakoid stacking is mediated by specific interactions between LHCII proteins across the stromal gap. Beyond these observations, our study establishes the methodological foundation to explore the conservation of thylakoid molecular architecture with other ecologically important phototrophs and dissect the mechanisms that adapt this architecture to changing environmental conditions.

## METHODS

### Cell Culture

We used the *Chlamydomonas reinhardtii* strains *mat3-4* (CC-3994) *(41)* and wild-type CC-125, provided by the *Chlamydomonas* Resource Center, University Minnesota, MN. The *mat3-4* strain has smaller cells that vitrify better by plunge freezing. Comparative 77K fluorescence measurements showed that *mat3-4* and wild-type cells have similar distributions of photo-synthetic antennas bound to PSI and PSII (Fig. S8). Both strains are in the same state, which is close to state I *(10)*. Thus, our description of the arrangement of photosynthetic complexes within *mat3-4* thylakoids is likely comparable to wild-type thylakoids.

For cryo-ET, *mat3-4* cells were grown to mid-log phase (1000-2000 cells/μL) in Tris-acetate-phosphate (TAP) medium under constant light conditions (∼ 90 µmol photons m^-2^s^-1^) and bubbling with normal air. For 77K measurements, both *mat3-4* and wild-type strains were grown under these same conditions, and measurements were made both immediately and after allowing cultures to sit in low light for 30 minutes.

### 77K Measurements

Fresh colonies of CC-125 and CC-3944 were picked from agar plates and suspended in sterile air-bubbled flasks containing 50 mL TAP medium. The cultures were grown at room temperature under 90 µmol photons m^-2^s^-1^ light until reaching 1,000 cells/μL. Samples containing 15 mL culture suspension were transferred into duplicate 15 mL Falcon tubes. From one of these tubes, five 1 mL aliquots were transferred to Eppendorf tubes and frozen immediately in liquid nitrogen. The second Falcon tube was kept under dim light of ∼ 10 μmol photons m^-2^s^-1^ for 30 minutes, with gentle mix by inversion, and five 1 mL samples were then frozen as above. Fluorescence emission spectra were acquired as previously described *(42)* under liquid nitrogen, using a custom-made apparatus. Excitation was from a 440 nm diode laser, the output of which was sent through one branch of a bifurcated optical fiber (diameter of 1 mm) to the sample. The emission light was collected through the second fiber and measured with a spectrometer (Ocean Optics HR200 + ER). The average 77K profile for each condition was composed of three independent biological replicates, each with five technical replicates (15 measurements total per condition).

### Vitrification and Cryo-FIB Milling

Using a Vitrobot Mark 4 (FEI Thermo Fisher), 4 µL of *mat3-4* cell culture was blotted onto R2/1 carbon-coated 200-mesh copper EM grids (Quantifoil Micro Tools) and plunge frozen in a liquid ethane/propane mixture. Grids were stored in liquid nitrogen until used for FIB milling. Cryo-FIB milling was performed following a previously-reported procedure *(22, 43)*, using a Quanta dual-beam FIB/SEM instrument (FEI Thermo Fisher), equipped with a Quorum PP3010 preparation chamber. Grids were clipped into Autogrid support rings modified with a cut-out on one side (FEI Thermo Fisher). In the preparation chamber, the frozen grids were sputtered with a fine metallic platinum layer to make the sample conductive. Once loaded onto the dual-beam microscope’s cryo-stage, the grids were coated with a thicker layer of organometallic platinum using a gas injection system to protect the sample surface. After coating, the cells were milled with a gallium ion beam to produce ∼ 100 nm-thick lamellas (Fig. 1= 60-90 nm, Fig. S1A= 100-140 nm, Fig. S1C= 60-80 nm, Fig. S1E= 110-140 nm). During transfer out of the microscope, the finished lamellas were coated with another fine layer of metallic platinum to prevent detrimental charging effects during Volta phase plate cryo-ET imaging, as previously described *(22, 44)*.

### Cryo-ET

After FIB milling, grids were transferred into a 300 kV Titan Krios microscope (FEI Thermo Fisher), equipped with a Volta phase plate *(24)*, a post-column energy filter (Quantum, Gatan), and a direct detector camera (K2 summit, Gatan). Prior to tilt-series acquisition, the phase plate was conditioned to about 0.5π phase shift *(45)*. Using SerialEM software *(46)*, bidirectional tilt-series (separated at 0°) were acquired with 2° steps between −60° and +60°. Individual tilts were recorded in movie mode with 12 frames per second, at an object pixel size of 3.42 Å and a target defocus of −0.5 µm. The total accumulated dose for the tilt-series was kept below ∼ 100 e-/Å^2^. Each tomogram was acquired from a separate cell and thus is both a biological and technical replicate.

### Tomogram Reconstruction

Frame alignment was performed with K2Align (https://github. com/dtegunov/k2align). Using IMOD software *(47)*, tilt-series were aligned with patch tracking, and bin4 reconstructions (13.68 Å pixel size) were created by weighted back projection. Of the 13 Volta phase plate tomograms acquired, four tomograms (Figs. 1 and S1) were selected for analysis of photosynthetic complexes based on good IMOD tilt-series alignment scores and visual confirmation of well-resolved complexes at the thylakoid membranes.

### Membrane Segmentation

Segmentation of chloroplast membranes was performed in Amira software (FEI Thermo Fisher), aided by automated membrane detection from the TomoSegMemTV package *(48)*. Bin4 tomograms were processed with TomoSegMemTV to generate correlation volumes with high pixel intensity corresponding to membrane positions. The original bin4 tomograms and correlation volumes were imported into Amira, and the correlation volumes were segmented by 3D threshold-based selection, producing one-voxel-wide segmentations at the centers of the membranes. Using the 3D lasso selection tool, these segmentations were subdivided into appressed and non-appressed membrane regions (Figs. 1B and S1B, D, and F). To generate membranograms visualizing complexes directly protruding from the membrane surface (PSII, PSI, cyt*b*_*6*_*f*), the segmentations were grown by two voxels in all directions to produce a five-voxel-wide segmentation with a surface that matched the surface of the membrane. To generate membranograms visualizing complexes ∼ 10 nm above the membrane surface (ATP synthase, membrane-bound ribosomes), segmentations were grown by 10 voxels in all directions. To produce smooth 3D surfaces, the segmented voxels were transformed into a polygonal mesh with the “generate surface” command, decimated to 10% triangle density with the “remesh surface” command, and smoothed with the “smooth surface” command (50 iterations, 0.4 lambda). These surfaces were exported in the OBJ 3D model format.

### Membranograms and Particle Picking

Bin4 tomograms and corresponding membrane segmentations (OBJ models) were loaded into Membranorama software (https://github.com/dtegunov/membranorama). This software projects tomographic density onto the surface of a 3D membrane segmentation to create a membranogram. Segmentations that had been grown by two voxels in Amira had surfaces that intersected densities protruding directly from the membrane surface (PSII, PSI, cyt*b*_*6*_*f*). Segmentations that had been grown by 10 voxels in Amira had surfaces that intersected densities that were ∼ 10 nm above the membrane surface (ATP synthase, membrane-bound ribosomes). Importantly, the Membranorama software can dynamically grow and shrink 3D membrane segmentations in real time, enabling the user to interactively track how densities appear at different distances from the membrane surface (see Movie S2). The software also allows to-scale 3D models of each molecular complex (PDB: 5ZJI for PSI, 5XNL for PSII, IQ90 for cyt*b*_*6*_*f*, 6FKF for ATP synthase, 5MMM for ribosome) (35, 39, 49–51) to be mapped onto the membrane and compared to the tomographic densities. This is accomplished interactively by clicking the surface of the 3D membrane segmentation and using the mouse wheel to rotate each particle in the plane of the membrane. Using these features, we manually assigned membrane-associated densities to different classes of macromolecular complexes based on their positions relative to the membrane and their characteristic structural features (Fig. S2). For the luminal side of the membrane, we exclusively used segmentations that had been grown by two voxels. PSII was assigned to large dimeric densities projecting ∼ 4 nm from the membrane surface, and cyt*b*_*6*_*f* was assigned to small dimeric densities projecting ∼ 3 nm from the surface. For the stromal side of the membrane, we started with segmentations that had been grown by 10 voxels, assigning large round densities with ∼ 25 nm diameters to ribosomes and smaller round densities with ∼ 10 nm diameters to the F_1_ subunit of ATP synthase. The strator of ATP synthase was often observed as a small density adjacent to the larger F_1_ density (see Fig. S2B). After assigning the positions of these two complexes, we next loaded the corresponding segmentation that had been grown by two voxels and assigned PSI to small round densities projecting ∼ 3 nm from the surface that were not positioned directly under a ribosome or ATP synthase. This order of particle picking prevented mis-assignment of PSI to the stalk of ATP synthase or the translocon structures that attach ribosomes to thylakoid membranes. The clarity of membrane-associated densities varied between different membranes within the same tomogram, likely due to effects of the tomographic missing wedge on different membrane curvatures and orientations, as well as local differences in the quality of tilt-series alignment. Therefore, only larger complexes (ribosomes, ATP synthase, PSII) were assigned for membranes with lower-clarity densities. Of the 51 non-appressed membranes quantified in this study, all complexes were assigned in 28 membranes, all complexes except for cyt*b*_*6*_*f* were assigned in 2 membranes, only ATP synthase and ribosomes were assigned in 4 membranes, and only ribosomes were assigned in 17 membranes. All complexes were assigned in the 33 appressed membranes quantified in this study. Within appressed membranes, PSII and cyt*b*_*6*_*f* were assigned with high confidence. Within non-appressed membranes, ribosomes and ATP synthase were assigned with high confidence, whereas PSI and cyt*b*_*6*_*f* were assigned with lower confidence (denoted by an asterisk in Table 1).

### Subtomogram Averaging

We used subtomogram averaging as a structural confirmation of the membranogram-picked positions for PSII and ATP synthase (Fig. S3). Manually-assigned positions and orientations were exported from Membranorama and used as starting parameters for real space subtomogram alignment in PyTom software *(52)*. No classification was performed, and all membranogram-picked subvolumes were included in the averages (396 PSII particles, 639 ATP synthase particles).

### Analysis of Protein Complex Organization

#### Nearest neighbor distances within the plane of a membrane (Fig. S4)

Segmentations of essentially flat membrane regions were exported from Amira (FEI Thermo Fisher) as MRC volumes. Coordinates of the particles (PSII, cyt*b*_*6*_*f*, ATP synthase, ribosomes) assigned on each membrane region were exported from Membranorama software. Each membrane region was projected with its corresponding particles onto a flat plane to generate a 2D surface. Nearest neighbor distances between the particles were then measured using Matlab scripts calculating the shortest path between objects. Clustered poly-ribosomes exclude large regions of the stromal surface (see Fig. 2C-D), which can cause misleading nearest-neighbor measurements for ATP synthase. To avoid this, the membranes were cropped to exclude regions containing poly-ribosome clusters before measuring ATP synthase distances.

#### Overlap between adjacent membranes using EM densities (Fig. 4E-F)

To calculate overlap between EM densities from two adjacent membranes, 11 membrane pairs separated by the stromal gap and 6 membrane pairs separated by the thylakoid lumen were segmented in Amira and imported into Membranorama software. Using the tools in Membranorama, surfaces from each membrane were selected and overlaid along a vector orthogonal to both membrane surfaces, ensuring a geometrically accurate superposition of the membrane densities. Images of the overlaid membranograms were then analyzed in Fiji software *(53)* as described in Fig. S6. Thresholding and cropping the membranograms was necessary to decrease noise and avoid edge effects, respectively.

#### Overlap between adjacent membranes using membrane models (Fig. 4C-F)

LHCII light-harvesting antennas are poorly visualized by cryo-ET because they are almost entirely embedded within the thylakoid membrane. Nevertheless, we incorporated hypothetical PSII-associated LHCII complexes into our analysis by using the PSII core positions and orientations visualized in membranograms to generate membrane models containing C_2_S_2_M_2_-type PSII-LHCII supercomplexes. First, 3D models of appressed membrane regions were exported from Amira (FEI Thermo Fisher). In the Membranorama software, we manually aligned a correctly-scaled 3D model of the C_2_S_2_M_2_-type PSII-LHCII supercomplex (PDB: 5XNL)*(35)* with the EM densities observed for each PSII core particle. Coordinates and orientations of all the particles were exported from Membranorama. The PSII-LHCII supercomplex structure was filtered to 30Å resolution and then mapped at the assigned positions and orientations into the 3D volume of the membrane segmentation, generating a 3D model of a one-voxel-thick membrane with embedded PSII-LHCII supercomplexes. To compare these experimentally-determined models with simulated models containing randomly distributed supercomplexes, the same number of PSII-LHCII supercomplexes were placed one-by-one at random positions in the membrane (randomly-selected voxels of the one-voxel-thick 3D membrane segmentation), and each time rotated by a random in-plane angle before placing the next supercomplex particle. Whenever a newly placed particle overlapped with a preexisting particle, rotation of the new particle was first attempted to avoid overlap, and if this failed, the particle was moved to a new random position (see Fig. S7 for a 2D schematic representation of this 3D procedure). 100 random models were generated for each appressed membrane region.

To measure the relative membrane area covered by the placed supercomplex structures (Fig. 4E), the one-voxel-thick membrane segmentation was masked where it was intersected by the 3D structure volumes (PSII core alone, or the full C_2_S_2_M_2_-type PSII-LHCII supercomplex). This masked “occupied area” was then divided by the total number of voxels in the membrane segmentation to yield a percentage. To calculate the overlap of “occupied area” between adjacent membranes (Fig. 4F), the PSII-LHCII supercomplex structure was subdivided into separate PSII (C2) and LHCII (S2M2) regions using Chimera software *(54)*. Next, volumes of the complexes in one membrane were extended along a vector orthogonal to both membrane surfaces until they also intersected the adjacent membrane. These extended complex densities were then used to mask “occupied area” on both membranes. Similar to the overlap calculation for EM density (detailed in Fig. S6), the percent of overlap was calculated as the number of membrane voxels that were masked by complexes from both membranes (equivalent to white in Fig. S6) divided by the sum of all masked voxels on the membrane (equivalent to white + green + magenta in Fig. S6). “White” overlap voxels masked by complexes from both membranes were only counted once in this calculation. The schematic models in Fig. 4C-D provide a 2D representation of this 3D analysis.

## AUTHOR CONTRIBUTIONS

M.S. froze cells and performed the cryo-FIB milling. M.S. and B.D.E. acquired and reconstructed tomograms. D.T. developed the Membranorama software. W.W. analyzed the data with assistance from B.D.E (segmentation) and S.A. (subtomogram averaging and analysis of protein complex organization). A.K. performed the 77K measurements. J.M.P. and W.B. enabled instrumentation and provided funding. B.D.E. provided funding and supervised the project. M.S, D.T., and B.D.E. conceived the study. W.W. and B.D.E. wrote the paper, with input from all authors.

## ACKNOWLEDGEMENTS

We thank Radostin Danev, Guenter Pfeifer, and Mattias Pöge for technical and computational assistance, Stefan Pfeffer for helpful discussions, and Karin Engel for critically reading the manuscript. This work was supported by a grant from the Deutsche Forschungsgemeinschaft to B.D.E. (EN 1194/1-1 as part of FOR 2092). AK was supported by the U.S. Department of Energy (DOE), Office of Science, Basic Energy Sciences (BES) under Award number DE-FG02-91ER20021. Additional funding was provided by the Max Planck Society.

---

**Figure S1.**
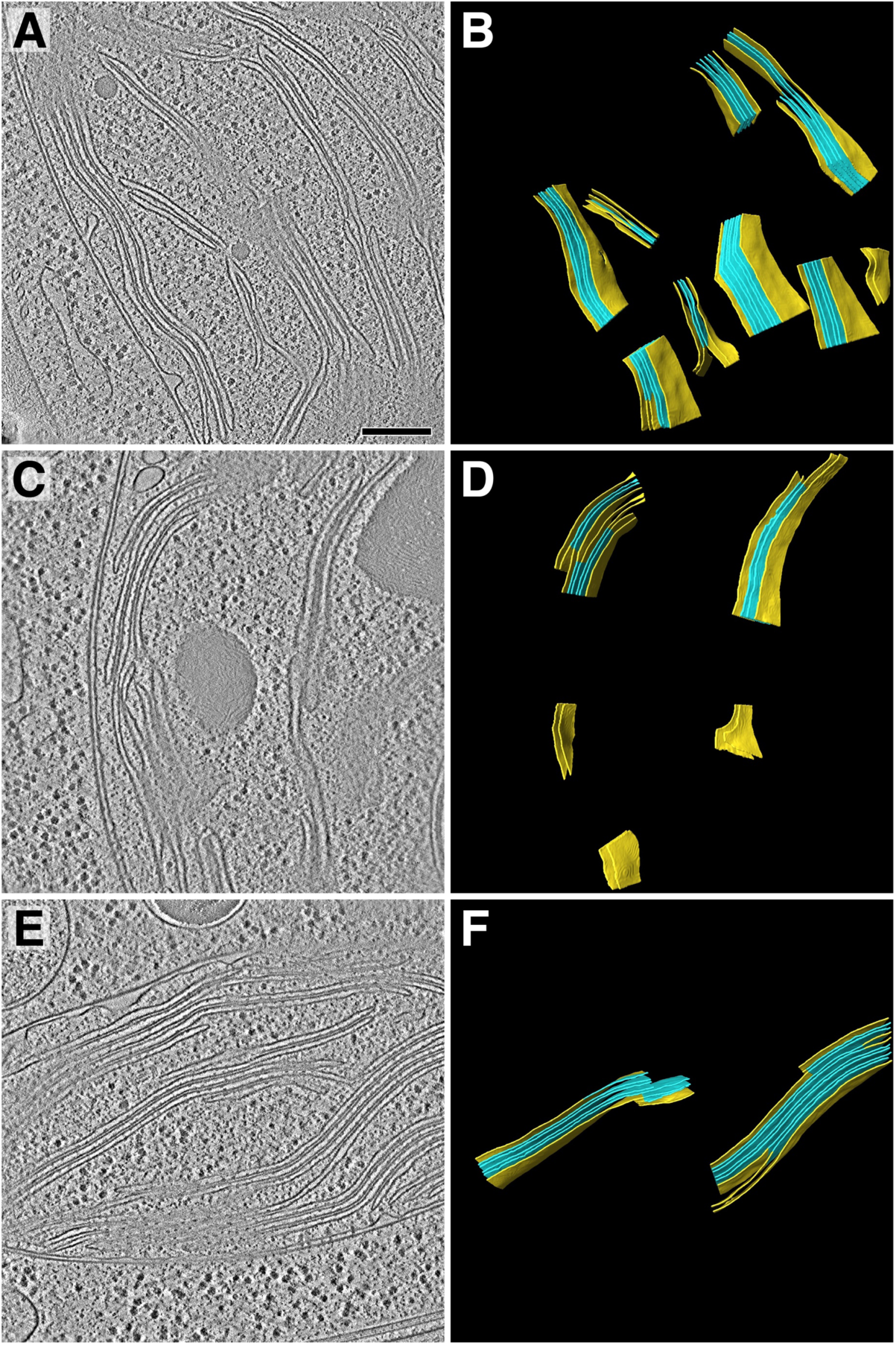
Tomogram overviews. Overviews of the three tomograms that were analyzed in addition to the tomogram shown in Fig. 1. Left panels show two-dimensional slices through the tomographic volumes, and right panels show three-dimensional segmented membrane regions (yellow: non-appressed, blue: appressed) where protein complexes were quantified by membranograms. Table 1 displays the total concentrations of protein complexes quantified from 84 membrane regions in the four tomograms. Scale bar: 200 nm.

**Figure S2.**
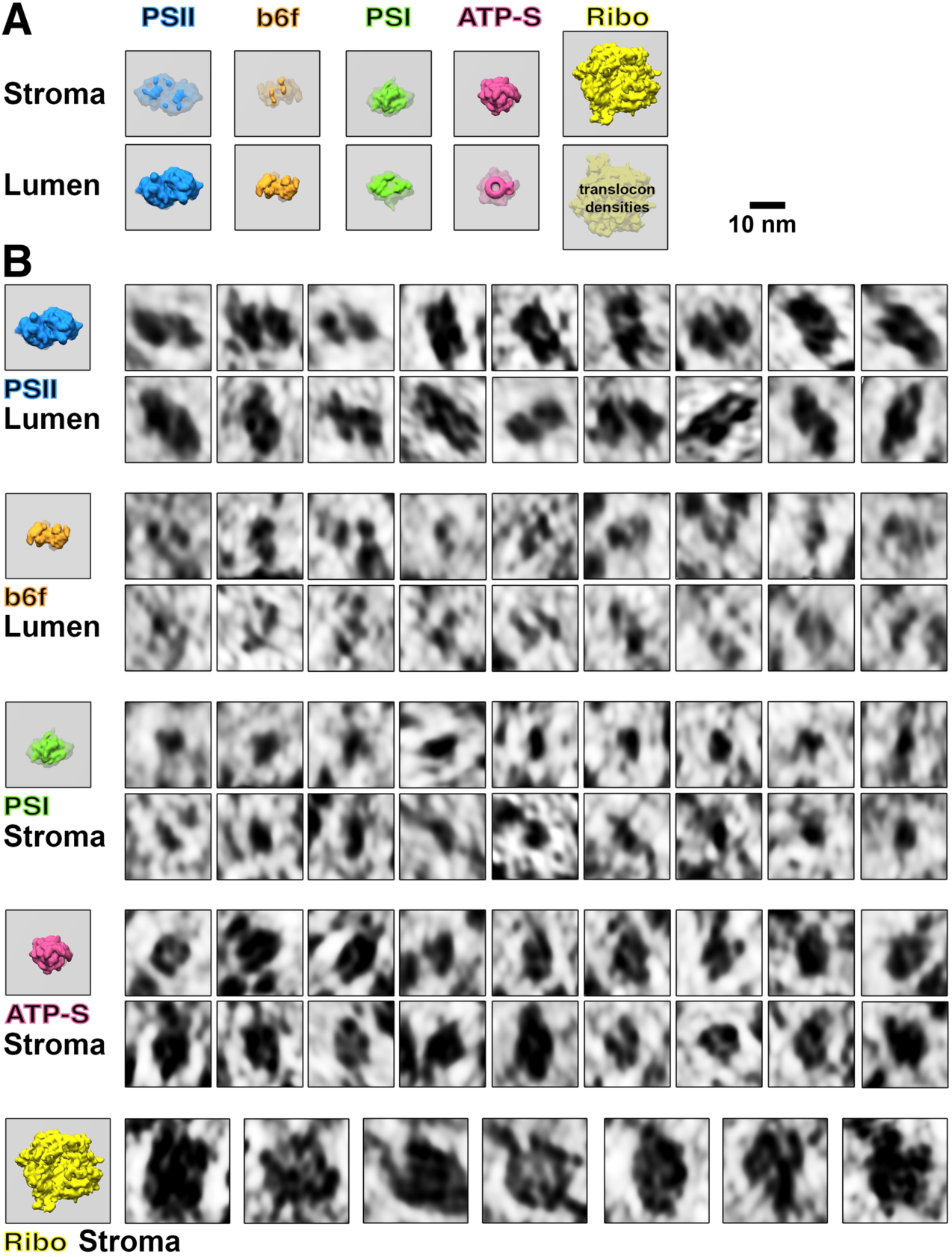
Membranogram particle gallery. **A)** Models of how each molecular complex appears when looking at the stromal and luminal surfaces of a thylakoid membrane (grey). Protruding densities are brightly colored, whereas densities located behind the membrane surface are greyer. Thylakoid-bound ribosomes would likely have small translocon densities visible on the luminal surface. Notice that outline “footprint” shapes of the complexes appear mirrored when viewed through the membrane. Photosystem II= PSII, cytochrome *b*_*6*_*f*= b6f, Photosystem I= PSI, ATP synthase= ATP-S, thylakoid-bound ribosomes= Ribo. **B)** Gallery of thylakoid-bound complexes as they appear in membranograms. Corresponding models of complexes protruding from the relevant membrane surface are shown on the left.

**Figure S3.**
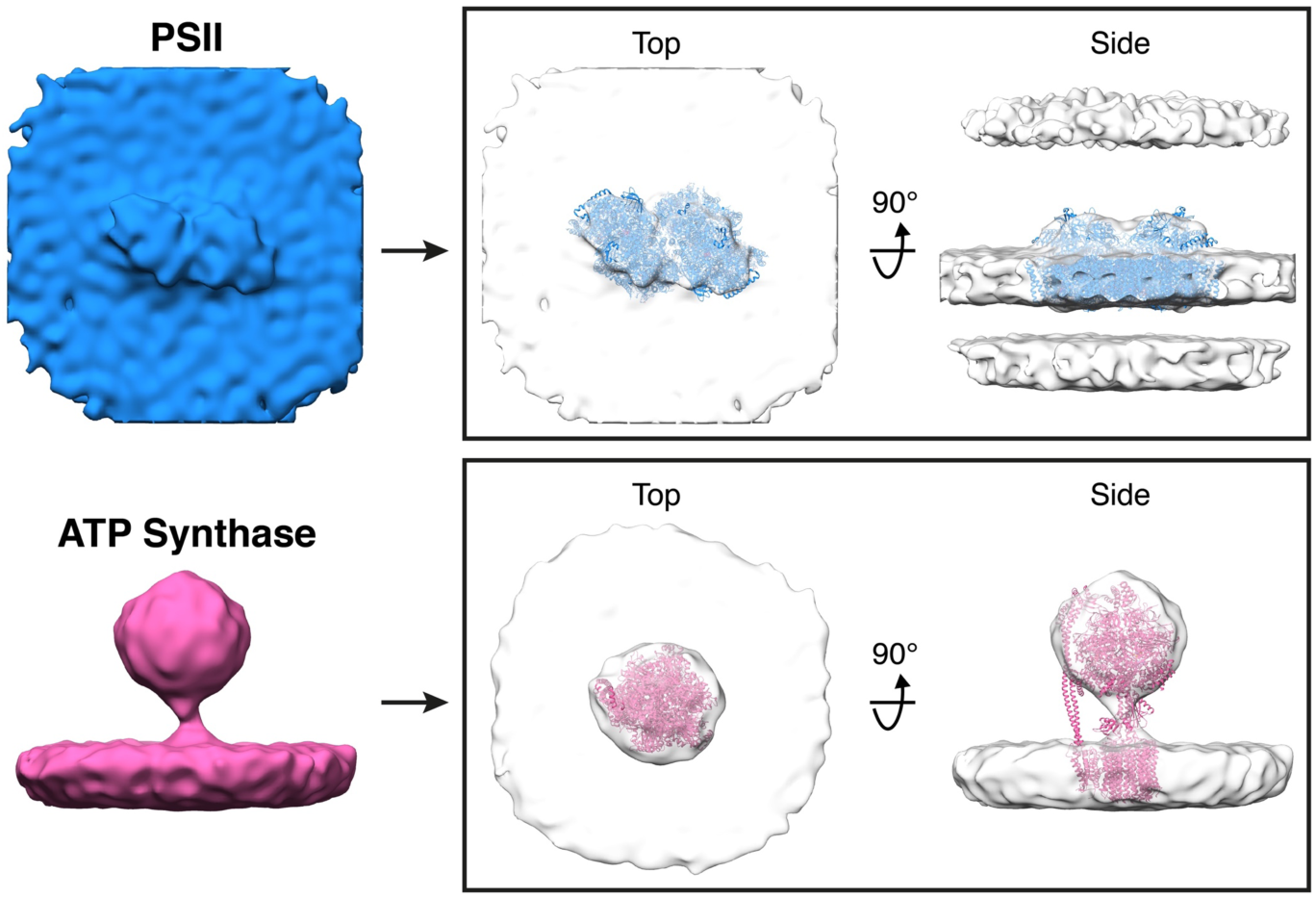
Subtomogram averages of PSII and ATP synthase generated from particle positions assigned by membranograms. In the boxes on the right, the averages are fitted with molecular structures determined by single particle cryo-EM (PDB: 5XNL, 6FKF) (*35, 39*). The subtomogram alignment did not resolve the rotationally asymmetric stator region of ATP synthase, likely because the F_1_ region undergoes large movements (*39*) that introduce structural heterogeneity in our active, native complexes. Nevertheless, both averages closely resemble the known structures of these complexes, validating the assignment of identities by membranograms.

**Figure S4.**
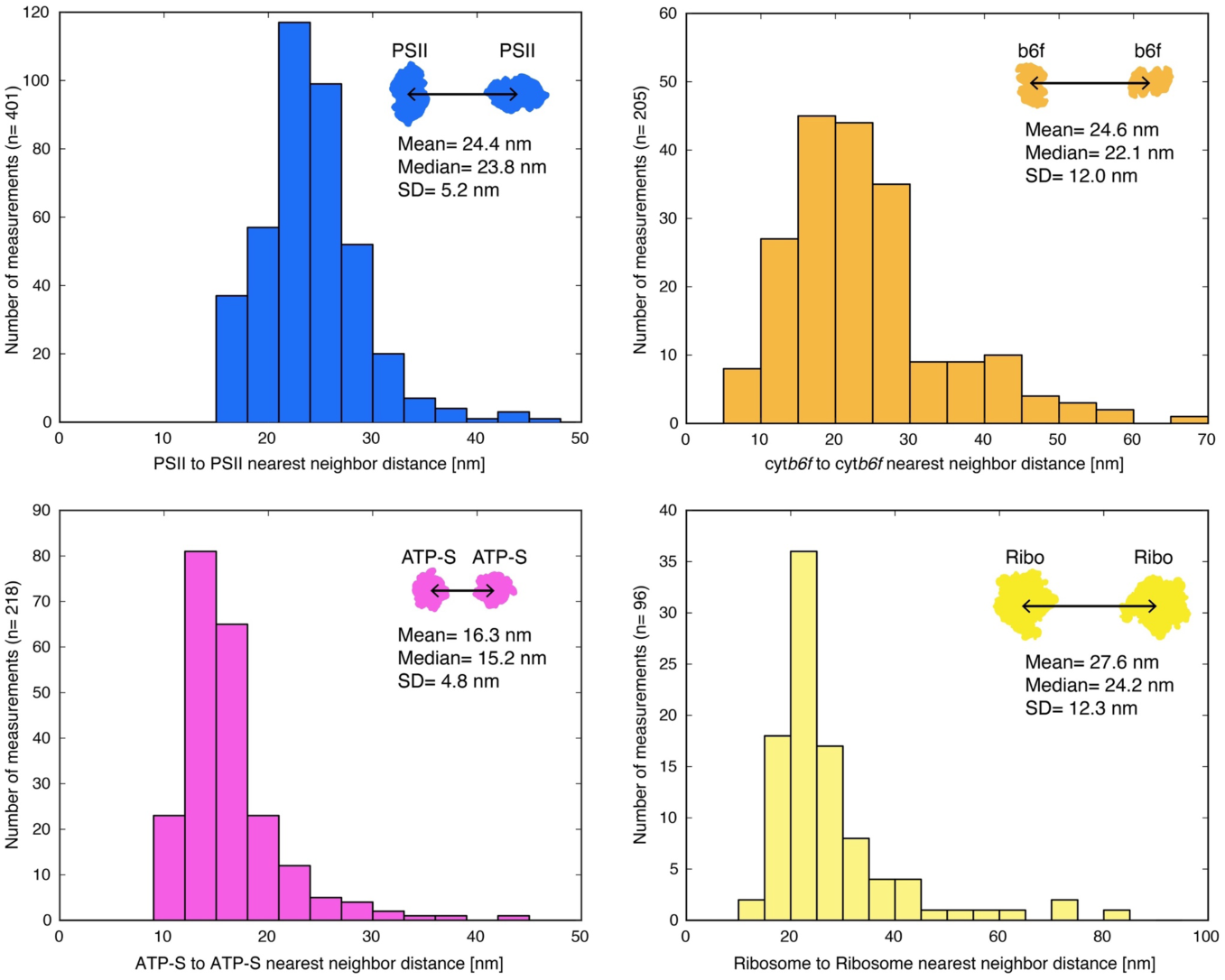
Distributions of nearest-neighbor distances for PSII, cyt*b*_*6*_*f*, ATP synthase, and membrane-associated ribosomes. Distances were measured between the center positions of nearest neighbor complexes within the plane of the membrane. Mean, median, and standard deviation of the mean (SD) are noted in the plots. The ribosome distribution has a sharp peak at 20-25 nm. This likely corresponds to the distance between ribosomes in thylakoid-bound polyribosome chains (Fig. 2C-D), as it is similar to the ∼ 22 nm ribosome spacing in bacterial polyribosomes (*40*).

**Figure S5.**
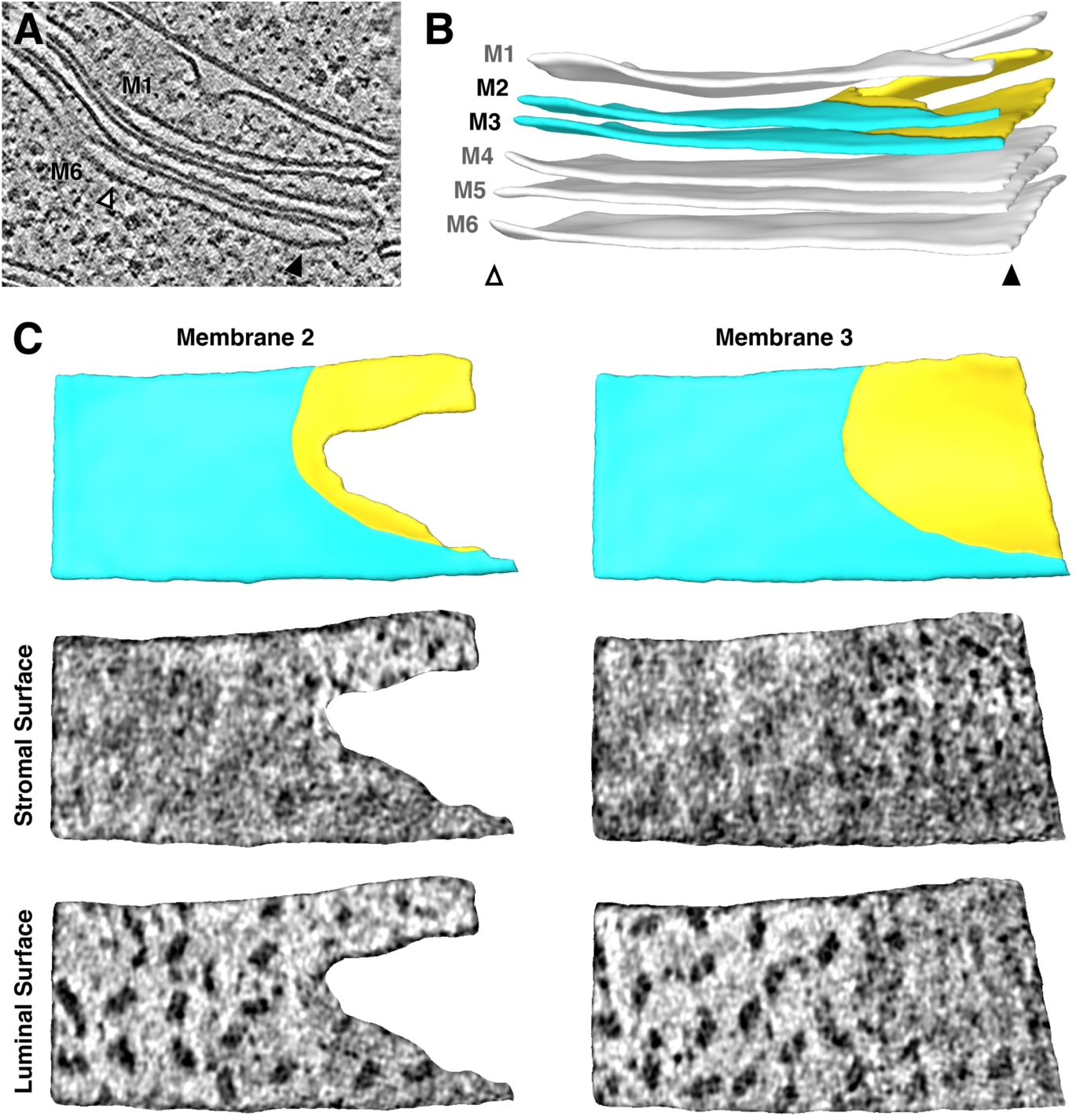
Additional example of the strict lateral heterogeneity between PSI and PSII at the transition between appressed and non-appressed membrane regions (Accompanies Fig. 3). **A)** Slice through a tomogram showing a stack of three thylakoids that splits into a stack of two thylakoids and a single thylakoid. **B)** Segmentation of the thylakoid membranes in this stack. Arrowheads and membrane labels (M1-M6) correspond to the regions indicated in **A**. The thylakoid composed of M1 and M2 is more architecturally complex than the region shown in Fig. 3. M2 and M3 (yellow: non-appressed, blue: appressed) are examined by membranograms. **C) Above:** Surface views of the segmented M2 and M3, showing the appressed and non-appressed regions of these adjacent membranes. **Below:** Membranograms of M2 and M3, showing densities ∼ 2 nm above the stromal and luminal surfaces. PSII is exclusively found in the appressed regions, whereas PSI is only seen in the non-appressed regions, with a sharp transition between regions.

**Figure S6.**
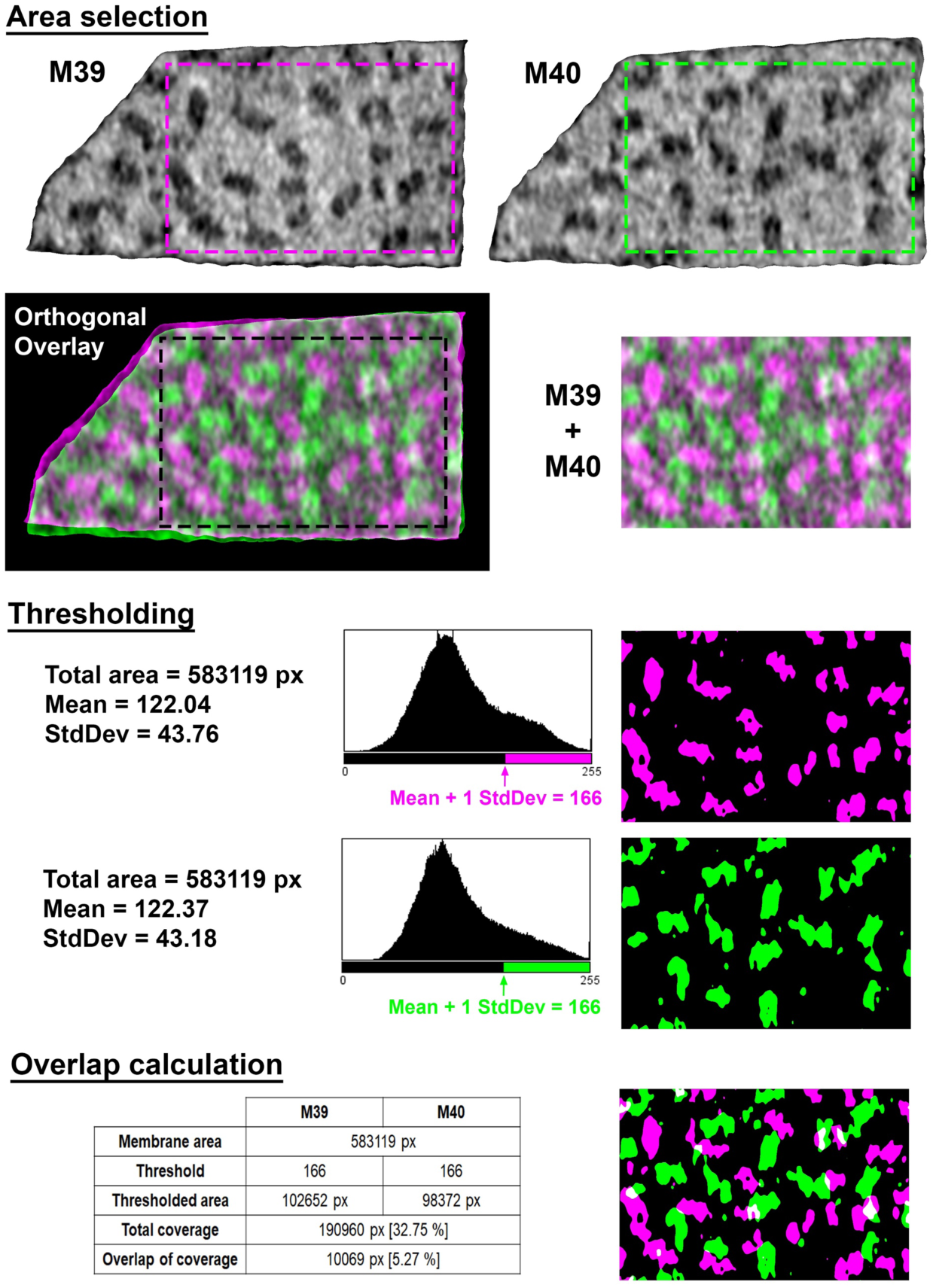
Calculations of membrane coverage and intermembrane overlap from EM density (accompanies Fig. 4). Area selection: To select overlapping areas of two adjacent membranes (M39 and M40 in this example; magenta and green, respectively), the two membranograms are overlaid along a vector orthogonal to both membrane surfaces. The overlay is then cropped to prevent edge effects (M39 + M40). **Thresholding:** Each cropped membranogram is binarized, with the threshold defined by the mean pixel value plus one standard deviation (StdDev), rounded to the nearest integer value. **Overlap calculation:** The percentage of each membrane occupied by EM densities was calculated as the number of thresholded pixels (green or magenta) divided by total pixels in the membrane (black + green or magenta). The percentage of overlapping EM density was calculated as the overlapping thresholded pixels (white) divided by the sum of the thresholded pixels in both membranes (white + green + magenta). Overlapping pixels were only counted once in this summation of all thresholded pixels.

**Figure S7.**
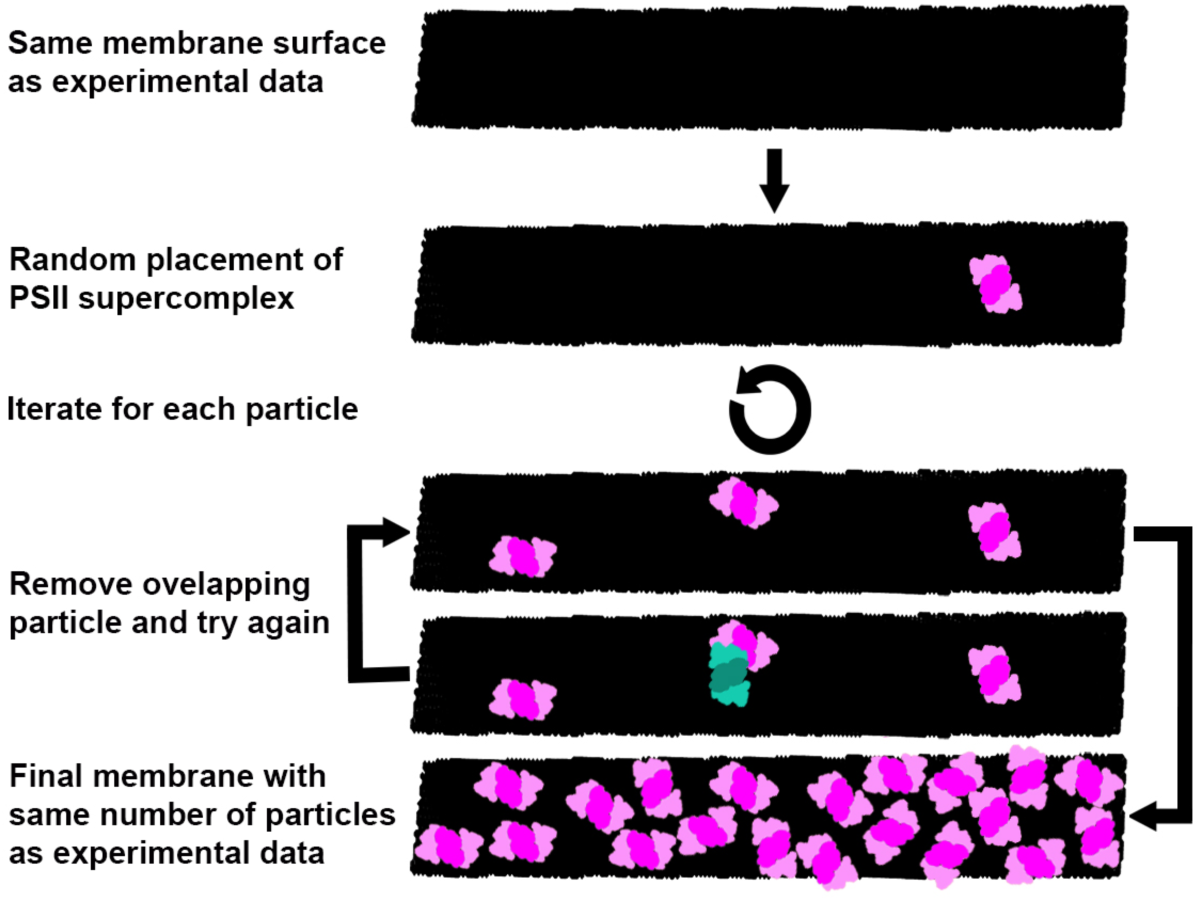
Generation of random membrane models (accompanies Fig. 4). Schematic representation of the workflow for generating membranes with randomly-distributed C_2_S_2_M_2_-type PSII-LHCII supercomplexes, used for the analysis of intermembrane PSII and LHCII overlap (Fig. 4F). The same number of supercomplexes (pink) as measured experimentally were placed one-by-one with random positions and orientations into the same membrane surface (black). If a new particle overlapped with a particle that had already been placed (event in blue/green), it was removed and placed again until it fit without overlap. The random distributions were simulated 100 times for each membrane. While diagrammed in 2D here for visual simplicity, the process was actually performed on 3D membranes (see methods).

**Figure S8.**
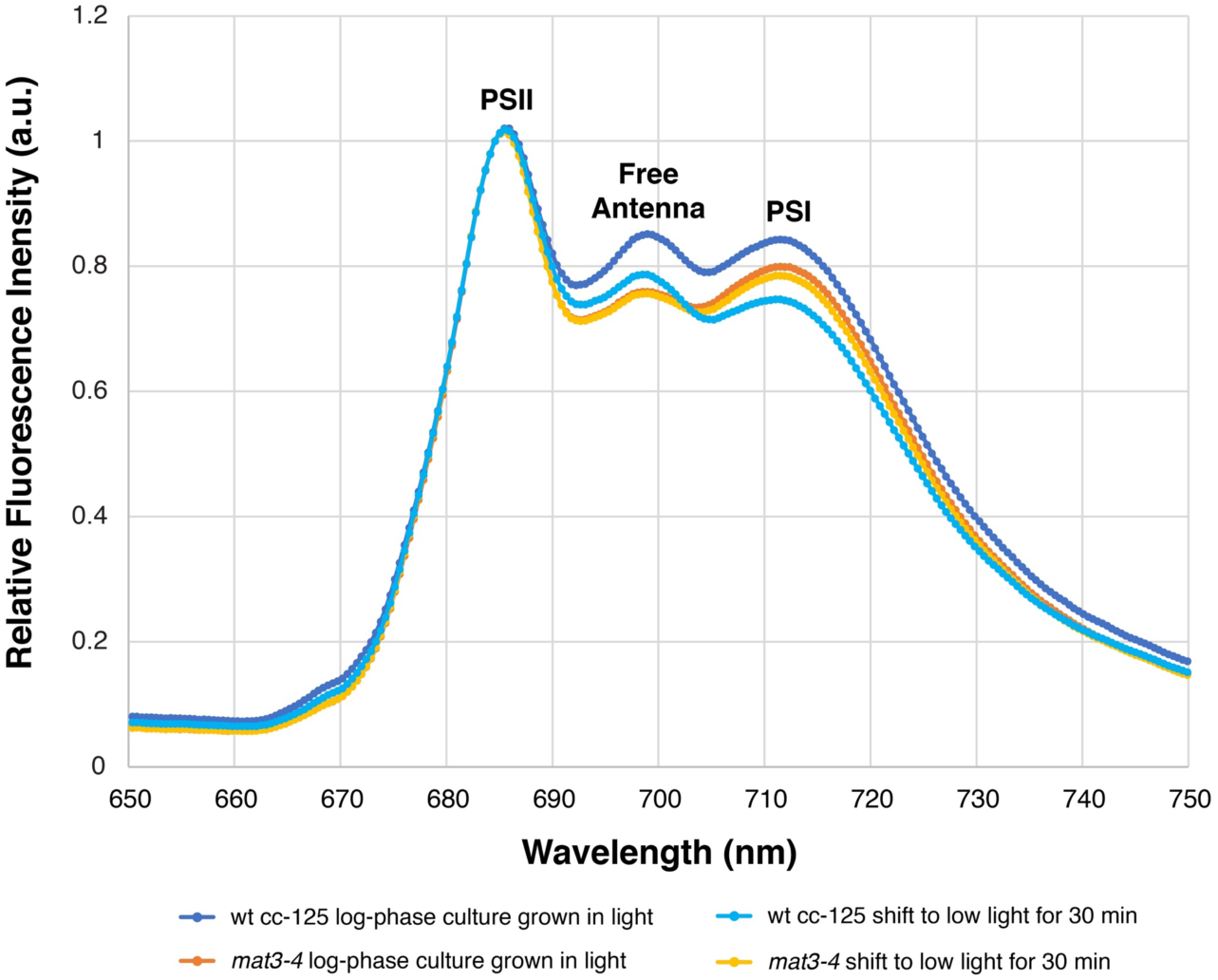
The *mat3-4* stain has a similar 77K fluorescence spectrum profile to wild-type cells. Measurements were made from log-phase cultures grown under the same conditions that were used for cryo-ET (illuminated with ∼ 90 µmol photons m^-2^s^-1^, bubbling with normal atmosphere). 77K spectra were measured from cells taken immediately from growing cultures (dark blue: wild-type cc-125, orange: *mat3-4*) and from cells that were allowed to sit for 30 minutes in a conical tube in low light (∼ 10 µmol photons m^-2^s^-1^), similar to the conditions that cells experienced before plunge-freezing. The plot of each condition is the average of three independent biological replicates, each with five technical replicates (15 measurements total per condition). These 77K spectra show that both *Chlamydomonas* strains in both environmental conditions were close to state I (*10*). Resting in a conical tube at low light without aeration did not induce anoxia or a shift cells towards state II. Since both strains have similar distributions of photosynthetic antennas bound to PSI and PSII, our cryo-ET description of the arrangement of photosynthetic complexes within *mat3-4* (Figs. 1-4; Table 1) is likely comparable to wild-type cells.

## Supplementary Movies

**Movie S1. In situ cryo-electron tomography reveals native thylakoid architecture with molecular clarity.** Sequential slices back and forth in Z through the tomographic volume shown in Fig. 1A. The yellow boxed region, focusing on the thylakoids shown in Figs. 2-4, is then enlarged and displayed in the lower right corner of the movie. Ribosomes and ATP synthase complexes can be seen bound to the stromal non-appressed thylakoid surfaces, while densities from PSII and cyt*b*_*6*_*f* extend from the appressed surfaces into the thylakoid lumen.

**Movie S2. Mapping molecular complexes onto the thylakoid architecture with membranograms.** Tomographic densities are projected onto the surface of a 3D membrane segmentation to produce a membranogram. The segmentation can be interactively grown and shrunk to visualize densities at different distances from the membrane. The bottom part of the movie shows a membranogram of the luminal surface of an appressed thylakoid membrane. In the top part of the movie, a moving red arrow indicates the position of the membranogram surface relative to the thylakoid architecture (shown both as real data and corresponding illustration). The movie begins by growing and shrinking the segmentation, showing densities that begin within the appressed membrane and extend into the thylakoid lumen. Growing and shrinking the segmentation immediately adjacent to the membrane surface allows careful inspection of the thylakoid-bound densities. First, the positions and orientations (boxes with vector lines) of the PSII and cyt*b*_*6*_*f* (b6f) complexes are assigned. To-scale 3D structures of both complexes are then mapped onto the membranogram, showing good overlap with the tomographic densities.

